# A new microcirculation culture method with a self-organized capillary network

**DOI:** 10.1101/2020.05.12.067165

**Authors:** Kei Sugihara, Yoshimi Yamaguchi, Shiori Usui, Yuji Nashimoto, Sanshiro Hanada, Etsuko Kiyokawa, Akiyoshi Uemura, Ryuji Yokokawa, Koichi Nishiyama, Takashi Miura

## Abstract

A lack of microcirculation has been one of the most significant obstacles for three-dimensional culture systems of organoids and embryonic tissues. Here, we developed a simple and reliable method to implement a perfusable capillary network in vitro. The method employed the self-organization of endothelial cells to generate a capillary network and a static pressure difference for culture medium circulation, which can be easily introduced to standard biological laboratories and enables long-term cultivation of vascular structures. Using this culture system, we perfused the lumen of the self-organized capillary network and observed a flow-induced vascular remodeling process, cell shape changes, and collective cell migration. We also observed an increase in cell proliferation around the synthetic vasculature induced by flow, indicating functional perfusion of the culture medium. We also reconstructed extravasation of tumor and inflammatory cells, and circulation inside spheroids including endothelial cells and human lung fibroblasts. In conclusion, this system is a promising tool to elucidate the mechanisms of various biological processes related to vascular flow.

## Introduction

Multicellular pattern formation has been one of the central issues in developmental biology [1‒3]. An extensively used tool to understand the mechanism of pattern formation is an organ culture system in which embryonic tissue is cultured at the air-liquid interface [4]. Recent advancements in stem cell biology have enabled the generation of small tissue structures from a single cell, which is called an organoid, and various organoids have been established [5].

A technical obstacle for standard culture systems of three-dimensional tissue structures is the lack of microcirculation. There are various methods to improve oxygen supply, such as culture inserts and rotator culture [6]. However, these methods cannot overcome the size limitation, i.e., if the cultured tissue size exceeds a 100-µm order, the tissue undergoes necrosis due to hypoxia [7]. Reproduction of a functional capillary network has not been successful. For example, the tube formation assay has been classically used to assess the pattern formation capability of endothelial cells [8], but the generated network structure lacks a functional lumen and is not perfusable.

Interactions between vascular endothelial cells and other types of cells are one of the main themes of study in vascular biology. A major example is pericytes that exist between basement membranes of endothelial cells, which stabilize the biological activities of the endothelial cells [9]. In addition, endothelial cells interact with circulating cells in blood [10]. For example, neutrophils transmigrate through endothelial cells at an inflammation site [11]. In cancer biology, hematogenous metastasis involves adhesion of tumor cells to endothelial cells and invasion [12]. However, an effective in vitro system to observe these phenomena is lacking.

In tissue engineering, various methods have been developed to implement a capillary network in a microfluidic device for perfusion in culture systems [13]. These methods are classified into two categories: predesigned and self-organization methods. Predesigned methods align endothelial cells by engineering techniques. Self-organization methods employ the spontaneous pattern formation capacity of cells to generate capillary network structures (reviewed in [6]). Recent advances in the integrative studies of tissue engineering and vascular biology have enabled construction of a perfusable vascular network in vitro. For example, in 2013, a microfluidic device was developed with a self-organized perfusable vascular network [14]. We have previously integrated a spheroid culture system with the self-organized capillary network to improve the culture conditions of spheroids [15, 16].

Currently, these new culture methods are highly technical and difficult to implement in a common biological laboratory. Various microfluidic chips are commercially available [17], but collaboration with engineering researchers is needed to obtain microfluidic devices with optimized designs. For perfusion itself, syringe pumps are not very common in a biological laboratory, and it is still technically difficult to connect tubes without collapsing the capillary network in a gel.

In the present study, we developed an easy method to enable microcirculation in ordinary glass-bottom culture dishes. First, we screened commercially available endothelial cells for their capacity to form a lumen. Next, we developed a culture system to perfuse culture medium in the lumen. The flow persisted for 12‒24 hours per one medium change, which enabled long-term perfusion. We observed the main features of the endothelial pattern formation, which correlated with flow, endothelial cell shape changes, collective migration towards the upstream of the flow, remodeling of the vascular network, extravasation of tumor cells, the effect of pericytes on pattern formation, and the perfusion of vascularized spheroids. These results show the usefulness of this culture method for elucidating various biological phenomena.

## Materials and Methods

### Cell culture

We used commercially available primary cultured cells to generate a perfusable vascular network based on a previous report [14]. The cells were human umbilical vein endothelial cells (HUVECs), human aorta endothelial cells (HAECs), human umbilical artery endothelial cells (HUAECs), human pulmonary artery endothelial cells (HPAECs), human microvasculature endothelial cells (HMVECs), and human lung fibroblasts (LFs) (Lonza Inc.). We used optimized growth media supplied by Lonza Inc. to maintain these cells. After screening for their pattern formation capability (Fig. S1), HUVECs were mainly used for experiments. We used LFs for the coculture system [18], which were maintained using FGM-2 culture medium and protocols provided by the manufacturer (Lonza Inc.). For visualization purposes, we used red fluorescent protein (RFP)-labeled HUVECs and GFP-labeled pericytes from Angio-proteomie Inc. HL60 and NMuMG-Fucci cells were provided by the Riken Bioresource Research Center (RCB2813 and RCB0041, respectively). Colon Tumor 26 (C26) is a colon cancer cell line isolated from a BALB/c mouse treated with carcinogen *N*-nitroso-*N*-methylurethan [19]. C26 cells were injected into the spleen, and cells that metastasized to the liver were isolated. By repeating this injection-isolation cycle four times, LM4 cells were isolated. The details of LM4 cells will be described elsewhere (EK, manuscript in preparation). HL-60 and LM4-GFP cells were maintained in RPMI 1640 medium (Nacalai Tesque, Inc.) supplemented with 10% FBS and 1% penicillin-streptomycin.

### Culture dishes

We developed a culture dish to exert static pressure on the self-organized capillary system. This dish consisted of standard 35- or 60-mm glass-bottom tissue culture dishes with a glass separator. The 12 phi glass well part was left open (Fig. 1a). This type of dish was obtained from a local manufacturer or assembled using a glass coverslip and bioinert adhesive such as replisil (silicone).

**Figure 1:**
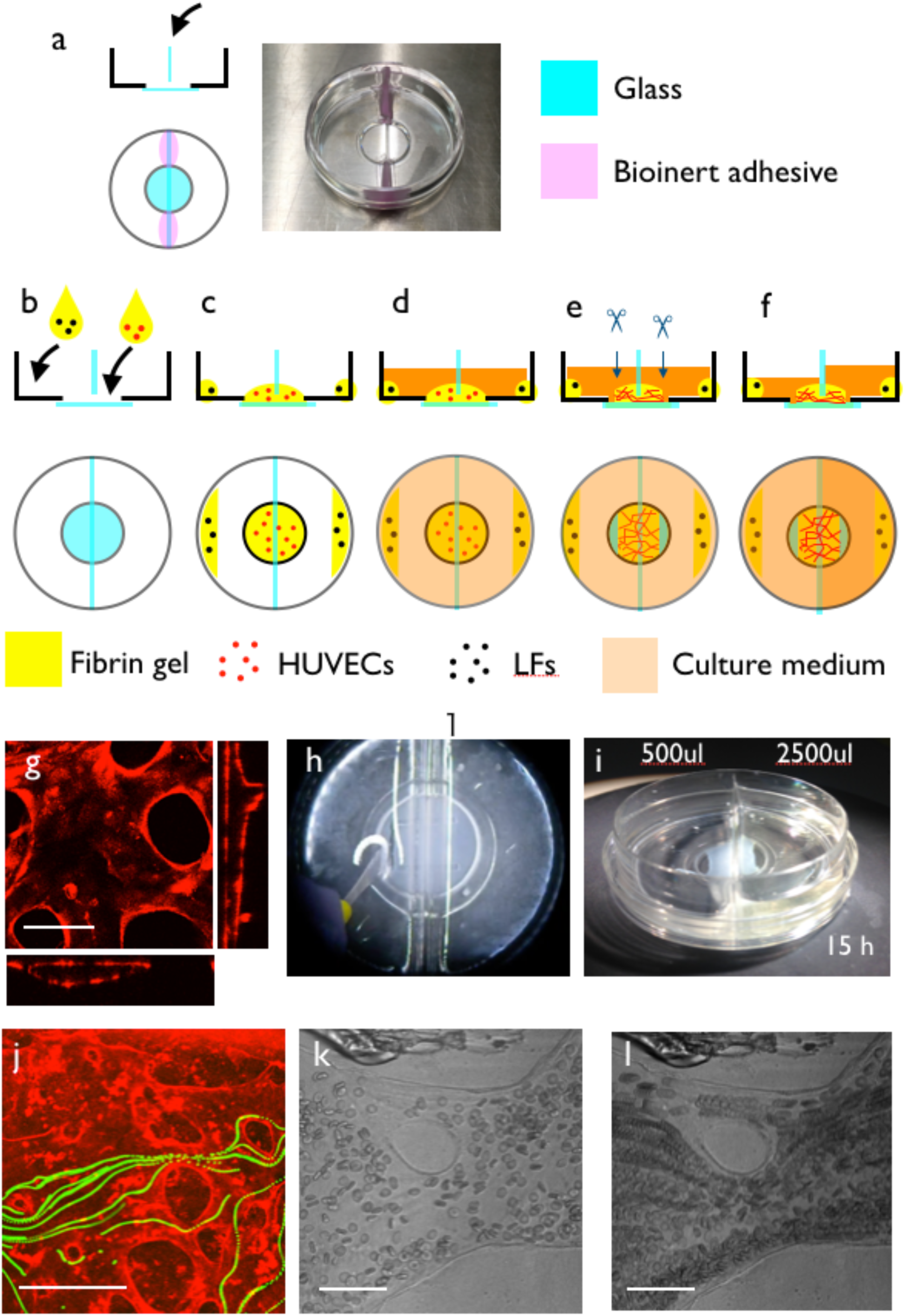
Culture system setup. (a) Side and top views of the culture dish before setup. The dish was a normal 12-phi glass-bottom dish. A glass separator was set at the center of the dish with a bioinert adhesive. (b) First, we placed 150 µl fibrin gel mixed with HUVECs in the center of the glass plate, so that two sides of the dish were separated by the glass separator and fibrin gel. We also added LF-containing fibrin gel at the edge of the dish. (c) We incubated the dish for 30 min to solidify the fibrin gel. (d) We added 1 ml culture medium to both wells and incubated the dish for 1 week. (e) After 1 week of culture, a vascular network with a perfusable lumen was formed in the glass-bottom region. Then, we cut both edges of the regions to make openings. (f) After the cuts, we increased the amount of culture medium on one side of the dish. This caused a static pressure difference between both openings of the self-organized capillary network, resulting in steady flow inside the apparatus. (g) A vascular network with a lumen was generated in the fibrin gel region. RFP-HUVECs were cultivated in the fibrin gel, and we observed the vascular network with lumens. (h) After the vascular network was formed spontaneously, both sides of the network were cut to make openings. (i) After making the cuts, we moved the culture medium, so we could exert static pressure to one side of the vascular network. The water level difference was maintained overnight, depending on the degree of lumen formation in the vascular network). (j) Flow inside the lumen was visualized using fluorescent beads. (k) Snapshot of the culture system when using whole blood as a tracer. Red blood cells were flowing inside the self-organized vasculature. (l) Projection of multiple frames of (k). Movements of red blood cells were visualized as a stream. Scale bars: 100 µm (g); 500 µm (j); 50 µm (k, l).

### Generation of a self-organized vascular network

In the first protocol, endothelial cells generated a capillary network by self-organization and the culture medium was perfused into the network by static pressure. First, we mixed 1 × 10^7^ HUVECs in a fibrin/collagen gel solution (5 mg/ml fibrin, 0.2 mg/ml type I collagen (rat tail, Enzo Life Sciences Inc., Alx-522-435-0020), and 0.15 U/ml aprotinin with a 1/150 volume of 0.5 U/ml thrombin). Then, we poured 150 µl of the fibrin/collagen/HUVEC solution into the well to separate the left and right halves of the dish divided by a glass separator. The dishes were incubated for 5 min at room temperature and then for 1 h at 37 °C to solidify the fibrin gel. Then, we added 5 × 10^5^ LFs in a 5 mg/ml fibrin gel solution at the edge of a 35-mm culture dish to avoid physical contact of LFs and HUVECs.

The dishes were incubated again for 1 h at 37 °C to solidify the fibrin gel. Then, we added 1 ml EGM-2 with 10 µM tranexamic acid to each well of the culture dish (total amount: 2 ml) and incubated the dish for 7 days (depending on the HUVEC activity, the culture period may be shorter). Culture media were changed once every 2‒3 days. After confirmation of capillary network formation with lumens, we cut the edge of the preformed network using tungsten needles or a sharp scalpel to make open ends. Then, we removed the culture medium from both wells and added 1 ml of culture medium to one of the wells. The existence of flow was confirmed by the flow of cell debris (Supplemental movie S2). Depending on the flow resistance, the water level difference persisted for 24 hours.

### Observation of vascular extravasation of neutrophils and tumor cells

Neutrophil-like differentiated HL-60 (dHL-60) cells were prepared by treating HL-60 cells with 1.25% DMSO (Nacalai Tesque, Inc.) for 2 days. Two pieces of thin PDMS sheet of roughly 6‒8 mm in size were placed on the glass bottom in parallel to each other and perpendicular to the glass separator before applying the HUVEC-containing fibrin gel. HL-60 and dHL-60 cells were visualized with CellTracker Green CMFDA (1:1000, Thermo Fisher Scientific). Then, 4 × 10^6^ HL-60, 1 × 10^6^ dHL-60, or 1 × 10^6^ LM4-GFP cells were suspended in 2 mL EGM-2 and applied to one side of the dish. For endothelial activation, 10 ng/mL human recombinant TNF-α (PeproTech, Inc.) was added to the medium overnight before the experiment. The HUVEC vascular network was visualized by incubation with rhodamine-conjugated UEA-I lectin (1:1000, Vector Laboratories) for 60 minutes in advance. Time-lapse and z-stack observations were performed with a Nikon A1R microscope.

### Generation of perfusable spheroids with vasculature

We also developed a protocol to perfuse spheroids with an endothelial cell vasculature (Fig. 8a). First, we prepared spheroids containing target and endothelial cells, and embedded the spheroid in a 2-well dish with fibrin gel. After the spheroids sprouted, we cut the tip of the sprout with a sharp scalpel and exerted static pressure for perfusion. Detail of the protocol is described below.

First, we prepared HUVECs, target cells, and lung fibroblasts. Then, we mixed HUVECs, lung fibroblasts, and target cells in a Sumiron MS-9096U 96-well dish. The ratio of HUVECs:LFs:target cells was 40:10:4. We cultivated the spheroids for 2 days.

Next, we embedded spheroids in the culture dish. We prepared the culture dish, fibrin, and thrombin solution. The culture dish was cooled on ice. We collected the spheroids in a 1.5-ml Eppendorf tube and removed the supernatants. Next, we added 150 µl of the fibrin gel solution to the Eppendorf tube on ice. Then, we added 1 µl of the thrombin solution to the Eppendorf tube, quickly agitated the spheroids, and then transferred the solution to the 2-well dish. We moved the dish to the stage of an inverted microscope (×2 objective lens) and carefully adjusted the locations of the spheroids using tungsten needles. The fibrin gel was solidified for 5 min at room temperature and then for 1 hour at 37 °C. During this period, we prepared LFs mixed with the fibrin gel solution. Finally, we added the thrombin solution to the LF-fibrin solution and then transferred the LF-fibrin gel solution to the side of the 2-well dish. After the fibrin gel solidified for 1 hour at 37 °C, we added 2 ml EGM-2 with 10 µM tranexamic acid and incubated the dishes for 7 days for the sprouts to elongate. In some cases, we seeded HUVECs on the solidified fibrin gel to prevent leak. Culture media were changed once every 2‒3 days.

### Flow tracers

We used various flow and permeability tracers to observe the characteristics of flow. Whole blood from a volunteer was diluted 10-fold with EGM-2 culture medium to observe red blood cell perfusion of the capillary network. Fluorescent particles (5 µm, Duke Scientific) were used to visualize flow for particle image velocimetry (PIV). FITC-dextran (70 kDa) was used for the permeability experiment. We also used diluted milk (1/1000) as a tracer for brightfield imaging.

### Histology and immunohistochemistry

Cultured blood vessels and spheroids were fixed in 4% PFA (immunohistochemistry) or Bouin’s fixative (HE staining) overnight. Fixed specimens were dehydrated using 70% ethanol in situ, detached from the culture dish, and transferred to a glass bottle. The samples were further dehydrated in graded concentrations of ethanol. Then, the ethanol was substituted with xylene and paraffin. The paraffin block was cut at 10 µm thicknesses. For histology, these sections were stained with hematoxylin and eosin.

For immunohistochemistry and Tdt nick end labeling (TUNEL), sections were deparaffinized and stained using the protocols provided by the manufacturers. Antibodies against PDGFB (Abcam, ab23914), type IV collagen (LSL, LB-0445), and desmin (LabVision, MS-376-S0) were used. To detect apoptotic cells, we used an In situ Apoptosis Detection Kit (Takara Bio, MK500).

### Image acquisition and analysis

Observation of three-dimensional structures was conducted using a Nikon A1R confocal microscope. For long-term observation of the whole culture dish, we used a Keyence BZ-900 with a tiling function. All image analysis was performed using ImageJ [20] or Fiji [21]. To observe vessel wall movement in long-term culture, we used the “linear stack alignment with SIFT” plugin for registration of the figures obtained at various time points. We used the “Reslice” command to prepare kymographs of cell movements from time-lapse movies.

## Results

### Pattern formation capability of commercially available primary cells

First, we screened commercially available human endothelial cells for their pattern formation capacity. The tested cells were HUVECs, HAECs, HMVECs, and HPAECs (Fig. S1). These cells were cultivated in normal glass-bottom dishes, Matrigel, collagen gel, and fibrin gel. In principle, all tested primary cell cultures were capable of generating a network of perfusable lumens only when the cells were cultivated in fibrin gel with human LFs, indicating that we could choose any type of primary endothelial cell. One exception was HPAECs that generated a perfusable network in fibrin gel without lung fibroblasts. This characteristic was observed in cells with low passage numbers up to 3.

### Method for culture medium perfusion

Next, we developed a culture method to perfuse the self-organized network in a glass-bottom dish. Using the protocol described in the Materials & Methods, we induced flow in the self-organized capillary network. First, we prepared a glass-bottom dish separated by a glass separator (Fig. 1a). Next, we allowed the cells to self-organize and generate a vascular network with a lumen in a gel that separated two wells of the dish. A mixture of HUVECs and fibrin gel was placed in the well of the glass-bottom dish (Fig. 1b) and allowed to solidify for 30 min. Lung fibroblasts for coculture were mixed in fibrin gels and placed at the edge of the culture dish to avoid contamination of lung fibroblasts in endothelial cell networks because fibroblasts inhibited formation of the endothelial cell network by direct contact (Fig. 1c). Finally, the culture medium was added, and the cells were incubated in a CO_2_ chamber (Fig. 1d).

After 1 week, the vascular network with a lumen was generated spontaneously (Fig. 1e, g). Next, we exerted static pressure on the preformed capillary network as follows. First, we made a cut to generate open ends of the capillary network (Fig. 1e. h). Then, we removed the culture medium from both wells and added culture medium (and tracer) to one of the wells (Fig. 1f, i). If the lumens were connected, we observed flow using the various tracers. We perfused fluorescent beads to detect the perfusability of the capillary network (Fig. 1j, Supplemental movie S1). However, the fluorescent beads adhered to the endothelial cell surfaces and interfered with the fluorescence observation. Therefore, this method was used only once per experiment. When whole blood diluted with EGM-2 was loaded, we observed the flow of red blood cells in the self-organized capillary network (Fig. 1k, l, Supplemental movie S2).

### Formation of a lumen around a solid object

In this culture system, lumen formation only occurred near the periphery of the culture well. Thick, perfusable lumens were formed in the region at 1 mm around the culture well wall or separating glass plate. This was not due to the high cell density around the periphery of the dish because increased cell density did not induce lumen formation in the center of the dish (Fig. S2a‒d). We induced lumen formation by embedding hard objects in the gel. When we mixed glass beads (1 mm diameter) with HUVECs at the beginning of the culture and cultivated the cells for 120 hours, we observed lumen formation around the glass beads (Fig. S2e‒h).

### Flow-induced collective cell migration and cell shape changes in the culture system

In this system, we observed the dynamics of endothelial cells in response to flow (Fig. 2). It is known that, at a certain range of flow rate, endothelial cells migrate collectively toward the upstream of blood flow[22‒24]. We easily observed this phenomenon in our culture system. We used Hoechst 33342 as a vital stain of endothelial cell nuclei and exerted flow to a 35-mm culture dish. We then observed the collective upstream movement of endothelial cells (Fig. 2a‒c). In the region with flow, the cells migrated collectively upstream of the flow (Fig. 2b). The white arrow indicates the orbit of cell debris, indicating that the direction of cell movement was opposite to that of the flow. In contrast, only random movement of cells was observed in the non-flow region (Fig. 2c), indicating that this system is useful to assess flow-endothelial cell interactions.

**Figure 2:**
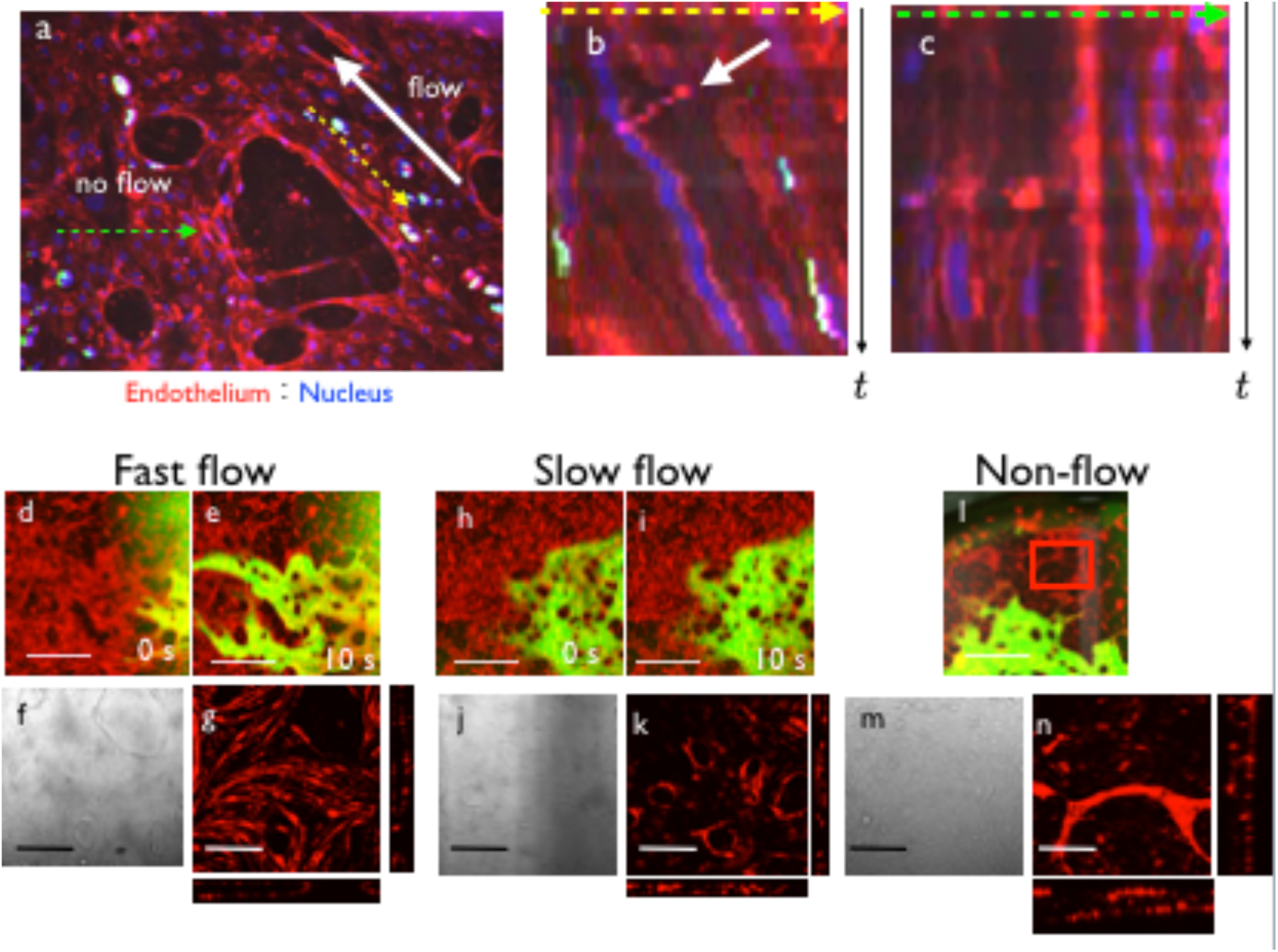
Flow-induced collective cell migration and cell shape changes. (a) Initial shape of the vascular network. HUVECs were stained with UEA1 and nuclei were stained with Hoechst 33342. There was a flow-positive region (yellow-dashed line) and non-flow region (green-dashed line). Direction of flow is indicated by a white allow. (b) Kymograph of the yellow-dashed line region. Collective movement toward upstream of the flow was observed. White arrow indicates the flow of cell debris. (c) Kymograph of the green-dashed line region. Cell movement was random. (d‒n) Cell shape changes induced by flow: (d, e) Fast flow. When FITC dextran was perfused, the vessel regions near the inlet or outlet showed fast flow. (f) Brightfield view of the fast flow region. Endothelial cells became shaped as spindles aligned parallel to the flow direction. (g) Fluorescence view of the fast flow region. At the floor of the lumen, we observed spindle-shaped cells parallel to the flow direction. (h, i) Slow flow. When FITC dextran was perfused, the vessel regions far from the inlet or outlet showed slow flow. (j) Brightfield view and (k) confocal view of the slow flow region. Endothelial cells did not show any polarity. (l) Low magnification view of the non-flow region. (m) Brightfield view and (n) confocal view of the non-flow region. Vasculatures were disconnected and became thin endothelial cysts with cell debris inside. Scale bars: 1 mm (d, e, h, i, l); 200 µm (f, g, j, k, m, n).

We next observed endothelial cell shape changes induced by the flow. In the fast flow region, endothelial cells became elongated parallel to the direction of flow (Fig. 2d‒g). In the slow flow region, the endothelial cells remained unpolarized (Fig. 2h‒k). In the non-flow region, the endothelial cells formed isolated cysts with cell debris inside (Fig. 2l‒n).

### Reconstruction of vascular remodeling by long-term flow in the vasculogenesis system

Because flow could be slow depending on the geometry of vasculature and the size of the cut, we maintained the flow for a long time. As the reservoir medium decreased, the pressure gradient changed. In general, we could maintain flow for 24 hours, which allowed long-term flow effects. We confirmed the flow by observing the flow of cell debris in the brightfield view (Supplemental movie S3). Unlike a two-dimensional culture system, HUVECs could be maintained for a very long period of up to 1 month without degradation of the fibrin gel (Fig. 3a‒c). The openings remained unobstructed after 4 weeks of culture (Fig. S3). A high magnification view revealed the occurrence of the remodeling process (Fig. 3d, e).

**Figure 3:**
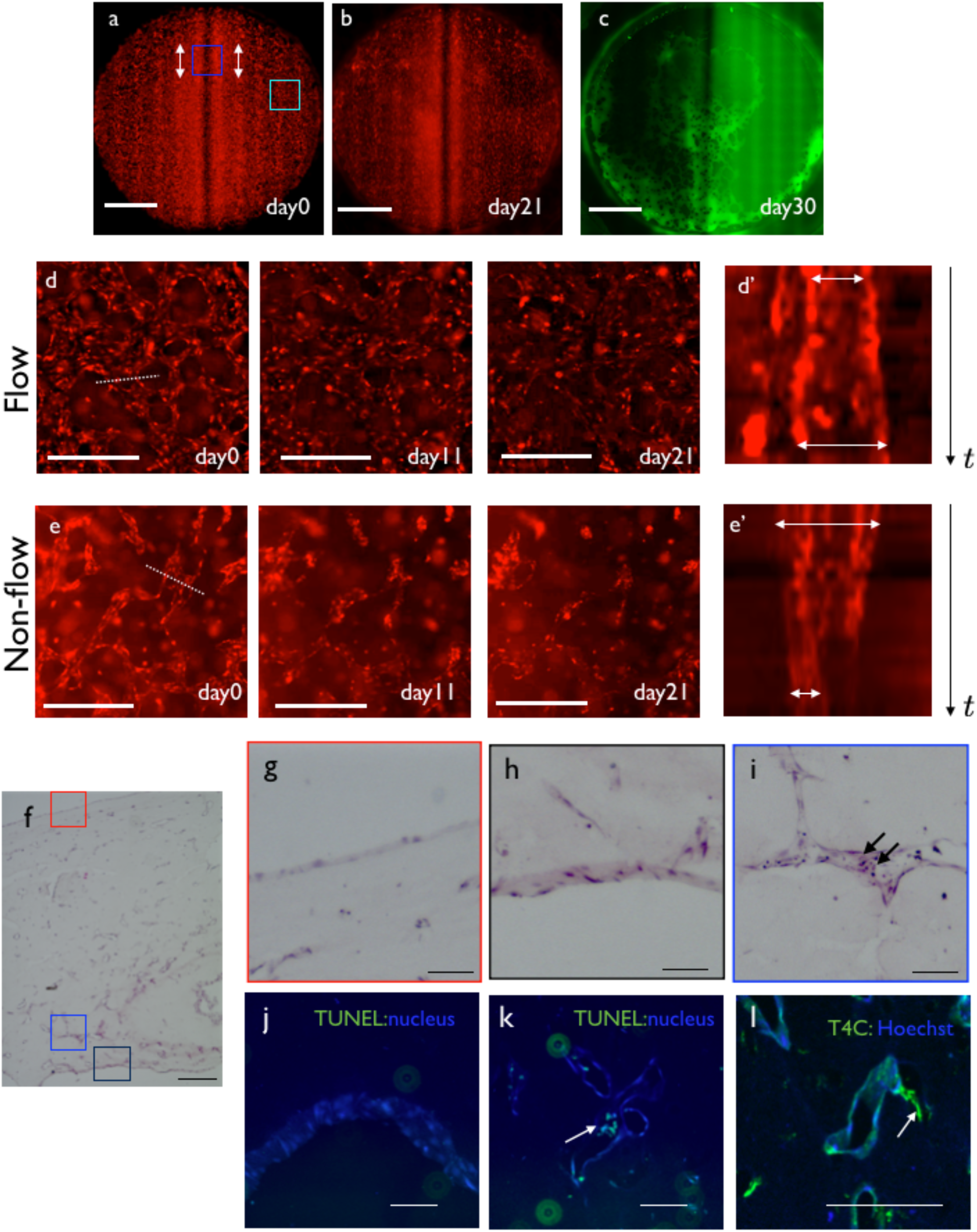
Reconstruction of flow-induced remodeling. (a‒c). Time course of the vascular network of the two-well dish. (a) Vascular pattern at day 0. Locations of the cut (white arrows) and high flow region (blue box), non-flow region (cyan box) are indicated. A cut was made at the periphery of the culture area to induce regions of flow and non-flow within a single dish. (b) Vascular pattern at day 21. (c) Visualization of the perfusable area by FITC-dextran. Perfusable regions existed near the inlet and outlet, near the glass separator and edge of the well. (d). Remodeling process at the high flow region from day 0 to 21. (d’) Kymograph of the dotted line region in (d). Vasculature dilated gradually. (e) Remodeling process at the non-flow region from day 0 to 21. (e’) Kymograph of the dotted line region in (e). The vascular diameter decreased. (f) Low magnification view of a hematoxylin-eosin-stained EC monoculture sample. (g) Magnified view of the upper surface of the gel. The surface of the gel was covered by endothelial cells. (h) Magnified view of the large lumen inside the gel. The lumen was also covered by endothelial cells. (i) Magnified view of the small lumen inside the gel. Small necrotic cells were observed inside the lumen (arrows). (j) TUNEL staining of the flow region. No dead cells were observed. (k) TUNEL staining of the non-flow region. Dead cells were observed inside the vasculature. (l) Type IV collagen staining of the long-term culture sample. There were some lumens positive for type IV collagen without cells (arrows), indicating the ECM sheath of the degraded vasculatures in non-flow regions. Scale bars: 3 mm (a‒c); 500 µm (d‒f); 100 µm (g‒l).

In the region with flow, the radius of the vascular segment was increased gradually (Fig. 3d, d’, Supplemental movie S4). In the region without flow, the vasculature was degraded gradually and finally became fragmented vascular cysts (Fig. 3e, e’, Supplemental movie S5).

Histological observation confirmed the remodeling process in this culture system. The top of the culture area was covered with HUVECs (Fig. 3f, g). The flow region was covered by a relatively thick HUVEC sheet (Fig. 3f, h). The non-flow region contained several apoptotic bodies (Fig. 3f, i). To confirm the distribution of cell death, we applied TUNEL staining. Positive signal was observed in the non-flow region (Fig. 3j, k), confirming the histological observation. We also observed an extracellular matrix sheath in which the basement membrane remained, while endothelial cells were retracted in the non-flow region (Fig. 3l). These observations supported that vascular remodeling occurred in this culture system.

### Effect of vascular flow on cell proliferation

Because the physiological role of vascular flow is the transfer of oxygen and nutrients, we examined the effect of flow on cell proliferation outside the blood vessel. We used Fucci-containing NMuMG cells, with which we could dynamically observe the cell cycle, to easily assess the effect of oxygen-nutrient transport by the flow on the mesenchyme region. NMuMG cells were mixed with RFP-HUVEC suspensions and cultivated for 1 week. The NMuMG cells formed sparse colonies in the region away from the vasculature, and the vasculature became narrower, presumably due to the competition between HUVECs and NMuMG cells (Fig. 4a). Next, we made a cut to only a part of the network to form flow and non-flow regions (Fig. 4b). We then compared the number of proliferating cells in flow and non-flow regions and detected increased fluorescence by exerting flow for 2 days (Fig. 4c‒e), indicating that the medium flow was effective for tissue maintenance in this culture system.

**Figure 4:**
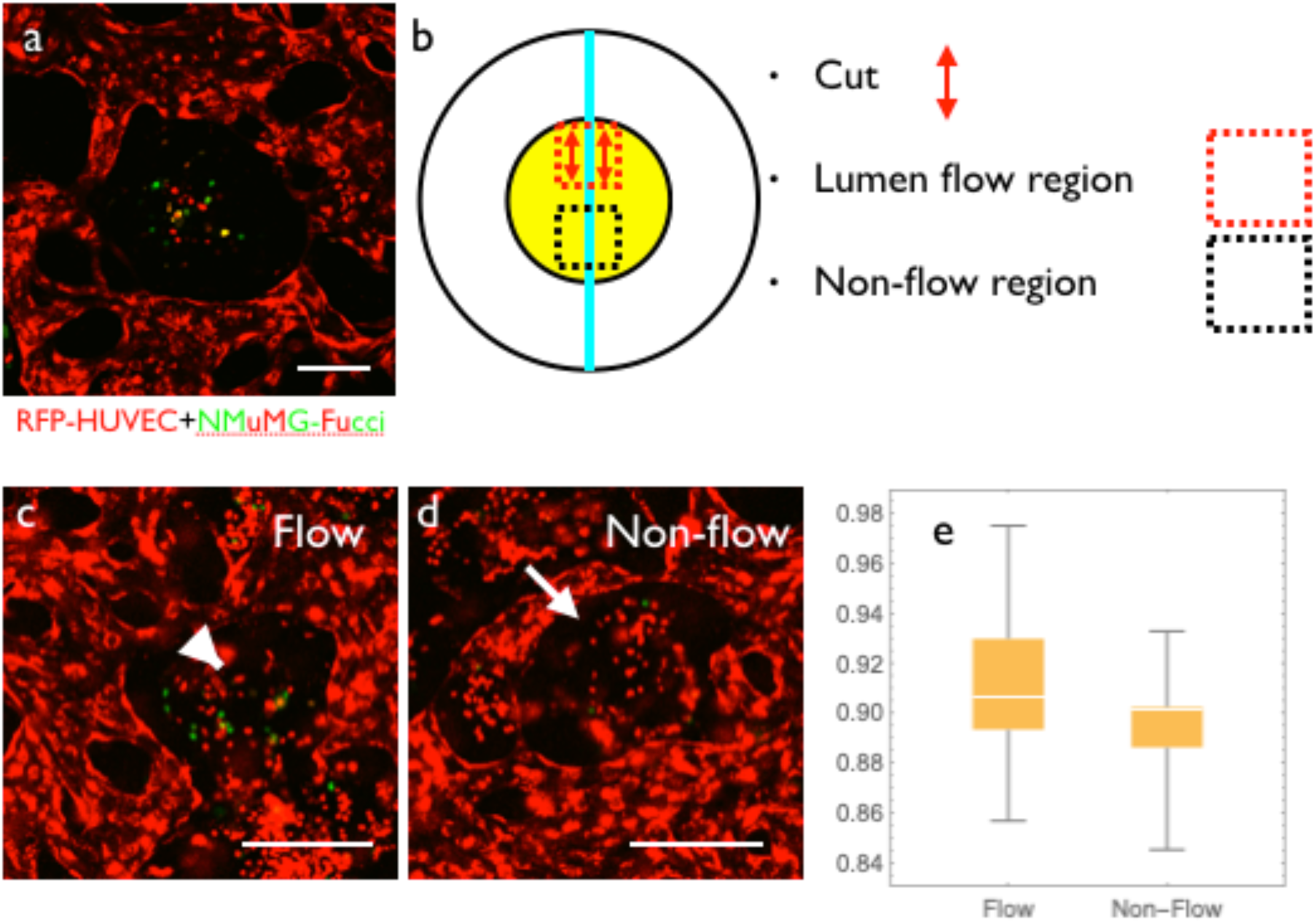
Effect of flow on cell proliferation. (a) Coculture of RFP-HUVECs and NMuMG-Fucci cells. HUVECs and NMuMG cells segregated, and NMuMG cells formed a colony in the interstitial region. (b) Experimental design. The upper region (red-dotted line) was cut to induce flow inside the lumen. We did not observe flow in the lower region (black-dotted line). (c) Flow region at 48 h. Proliferative cells remained in the interstitial region (white arrowheads). (d) Non-flow region at 48 h. Proliferating cells disappeared in the interstitial region (white arrows). (e) Relative green fluorescence intensity ratio (24 h/0 h). A statistically significant difference was found between flow and non-flow regions (Mann-Whitney test, p<0.05). Scale bars: 200 µm.

### Dynamics of pericytes in the self-organized vascular network

Pericytes reside around endothelial cells and modulate their biological functions [9]. It has been reported that pericytes play an important role in the vascular remodeling process to generate a hierarchical vascular tree structure in developing retina [25]. To examine whether addition of pericytes to this culture system affected the remodeling process, we cocultured pericytes in this culture system and observed the long-term pattern change with flow. Vasculature with pericytes grew normally, and we maintained the culture system with pericytes for up to 1 month (Fig. S4a, b). After 1 month, the vasculature was still perfusable without leak (Fig. S4c, d). The morphology of vasculatures with pericytes was similar to that without pericytes (Fig. S4c, d). In the region with flow, the diameter of the vascular segment was increased (Fig. S4e, e’), whereas in the region without flow, the vasculature was degraded gradually (Fig. S4f, f’). Unfortunately, the final shape of the vascular network with pericytes appeared more or less the same compared to that without pericytes (Fig. S4a‒d).

Interestingly, pericytes disappeared around the fast flow region and surrounded the endothelial vasculature in the non-flow region (Fig. 5a, b). Initially, pericytes were distributed evenly (Fig. 5a). After 3 weeks of culture, spots of high pericyte cell density appeared in non-flow regions (Fig. 5b, arrows). However, in the fast flow region, the pericytes had disappeared (Fig. 5b, red circle). We observed active migration away from the fast flow region (Fig. 5c. d, Supplemental movie S6). In the fast flow region, a very small number of pericytes was observed (Fig. 5e). At the interface between flow and non-flow regions, we observed an increase of pericytes only in the non-flow region (Fig. 5f). In the non-flow region, we observed pericytes wrapped around cysts of endothelial cells containing debris (Fig. 5g). To understand the reason for this uneven distribution, we observed the distribution of PDGFB, a chemoattractant for pericytes (Fig. 5h‒k). In the flow region, the PDGF signal was low, while in the non-flow region, strong signals were detected in endothelial cell cysts and pericytes (Fig. 5h‒k). This result suggested induction of PDGFB in the non-flow region was one of the reasons why pericytes only remained in the non-flow region.

**Figure 5:**
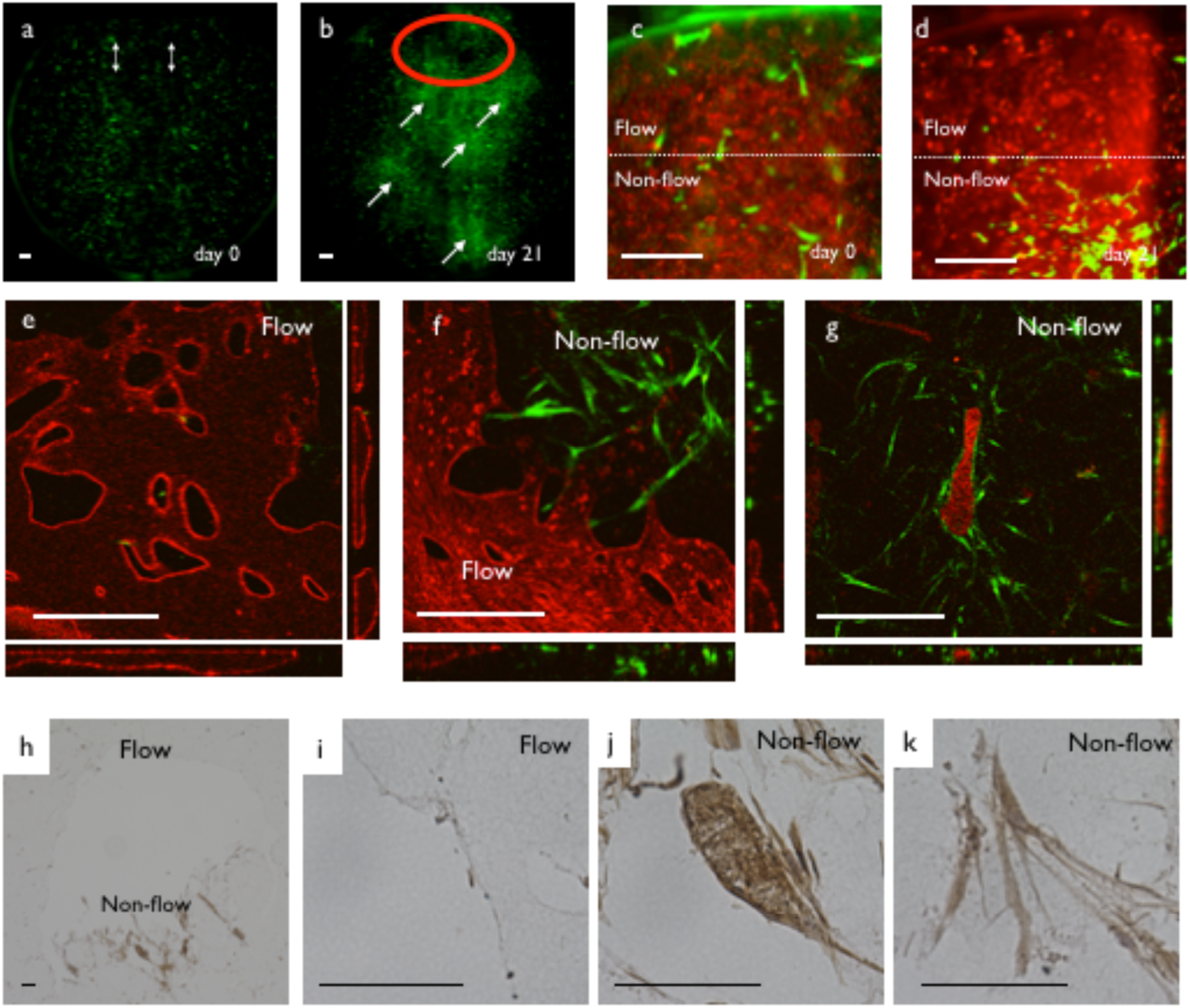
Dynamics of pericytes in the HUVEC-pericyte coculture system. (a) Low magnification view of the distribution of pericytes at day 0 (green). White arrows indicate the inlet and outlet. (b) Distribution of pericytes after 21 days of perfusion culture. We observed a high density region of pericytes (arrows) in the non-flow region and a low density region (red circle) in the flow region. (c) High magnification view of pericyte distribution at the flow region (day 0). (d) High magnification view of the pericyte distribution at the flow region (day 21). Pericyte cell density was decreased in flow region and increased in non-flow region. (e) 3D structure of the flow region at day 30. Endothelial cells (red) formed vasculature with a perfusable lumen. Pericytes were virtually absent. (f) Boundary between flow and non-flow regions. Pericytes resided in the non-flow region. (g) 3D structure of the non-flow region. Vasculature was degraded and flat cysts of endothelial cells with cell debris inside remained. Pericytes appeared to surround the degraded structure. (h) Low magnification view of PDGFBB immunohistochemistry. A positive signal was observed in the non-flow region. (i) High magnification view of the flow region. No positive signal was observed. (j) High magnification view of the non-flow region. The cell cyst and surrounding pericytes had a high staining intensity. (k) High magnification view of pericytes in the non-flow region. A positive signal was observed in the cytoplasm. Scale bars: 500 µm (a‒g); 100 µm (h‒k).

### Reproducing extravasation of inflammatory cells from the capillary network

We next observed the interaction of blood cells and capillaries using this synthetic system. At first, we perfused fluorescent beads (Fig. 1j) or red blood cells (Fig. 1k‒l) in the self-organized vascular network. We also perfused the neutrophil cell line HL-60 in the self-organized vascular network (Fig. 6a, b, Supplemental movie S). We differentiated HL-60 cells by DMSO treatment and perfused the cell suspension in the self-organized vascular network. The cells had the ability to move through a very thin capillary segment (Fig. 6c). We also observed the time course of extravasation (Fig. 6d‒f). Three-dimensional observation confirmed that the HL-60 cells were outside of the vascular network (Fig. 6g).

**Figure 6:**
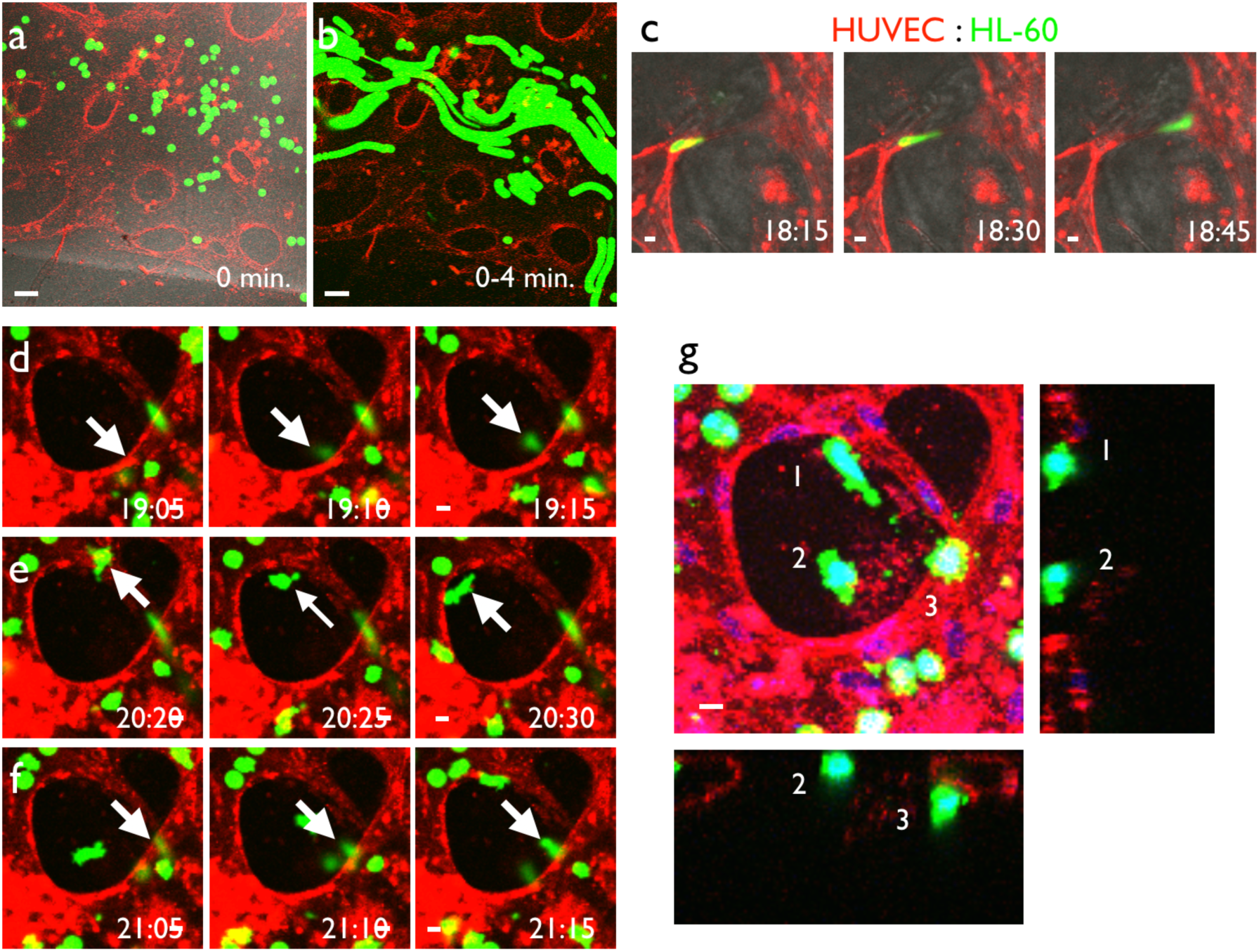
Extravasation of leukocytes using the self-organized vascular network. (a) HL-60 cells, an acute myelogenous leukemia cell line, were labeled with CellTracker Green and introduced into the self-organized vascular system visualized by UEA-I lectin. (b) Projection image of a time-lapse movie for 4 seconds. Movement of HL-60 cells inside the vascular network was observed. (c) DMSO-induced differentiated HL-60 cells (dHL-60), which mimicked neutrophils, passed through thin vessels with large deformation under the presence of TNFα. (d‒f) Representative time-course of extravasation. Extravasation occurred within a short time period (10 min.). Extravasating cells (white arrows) extended protrusions toward the outside of the blood vessel when emigrating from the blood vessel lumen. (g) Three-dimensional structure of extravasated cells (white arrows) shown as max projection x-y and x-z/y-z single slice images. We clearly observed extravasated cells outside of the vasculature. Time separated with colons indicates hours and minutes. Scale bars: 50 µm (a, b); 10 µm (c‒g).

### Reproducing metastasis of cancer cells from capillary vessels

The other possible application of this culture system is assessment of hematogenous metastasis. We introduced the cancer cell line LM-4 into the culture medium with perfusion and observed the dynamics for up to 48 hours. As a result, the cells flowed inside the self-organized vascular network (Fig. 7a). The cells also changed their shape during emigration from the vascular lumen (Fig. 7b). We next observed the detailed morphology of cancer cells after extravasation. The emigrated cells maintained adherence to the vascular wall after extravasation and extended protrusions along the endothelial cell surface (Fig. 7c-c’’), which may be regarded as filopodia-like protrusions in hematogenous metastasis [26]. In some cases, the extravasated cells influenced the shape of the vasculature (Fig. 7d, d’). We also observed vascular wall retraction at the sight of the extravasated cancer cells.

**Figure 7:**
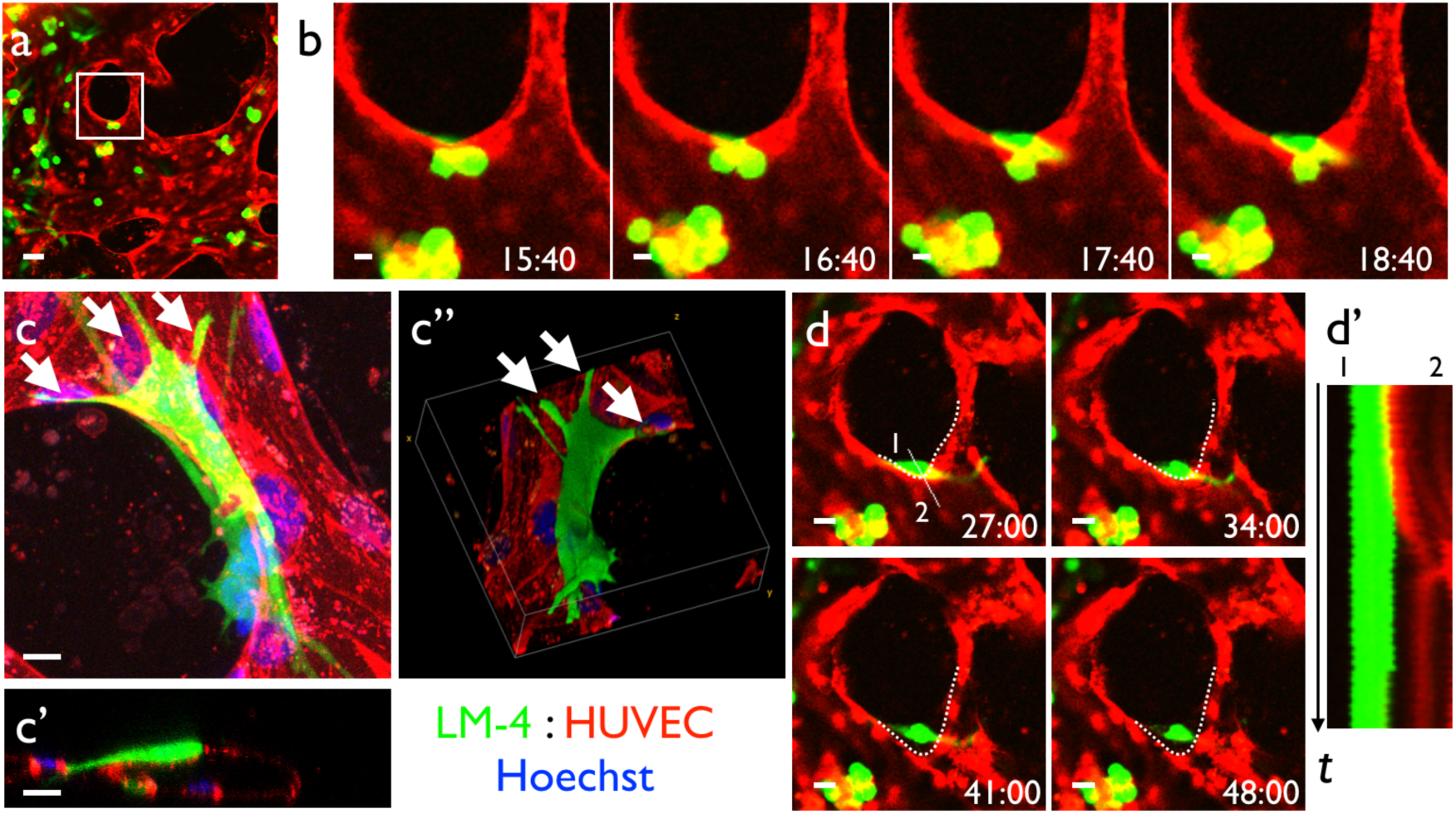
Reproduction of hematogenous metastasis of cancer cells from the capillary network. (a) LM4-GFP cells were introduced into the self-organized capillary network consisting of HUVECs visualized by UEA-1 lectin. (b) High magnification time-lapse view of (a). We observed cancer cells emigrating out of the blood vessels. (c) Detailed morphology of cancer cells on the endothelial cells. Emigrated LM4 cells attached to the blood vessel with highly polarized morphology and multiple protrusions. (c) Max projection image, (c’) orthogonal section, and (c’’) 3D-reconstructed image. White arrows: cancer cell protrusions. (d) Blood vessel shape changes after cancer cell emigration (white-dotted line). (d’) Kymograph of the blood vessel wall corresponding to the white line in (d). Time separated with colons indicates hours and minutes. Scale bars: 50 µm (a); 10 µm (b, c); 20 µm (d).

### Vascularized spheroid culture system with flow

We reproduced the spheroid culture system with perfusable vasculature [15] with a slight modification of the protocol (Fig. 8a). First, we produced spheroids using a 96-well plate. Then, the spheroids were embedded in the fibrin gel underneath a glass separator. When we cultured the spheroids for 1 week, HUVECs formed sprouts with a lumen, which were as long as 1 mm (Fig. S5). Then, we cut the tip of the sprouts to perfuse culture medium inside the vascular lumen.

**Figure 8:**
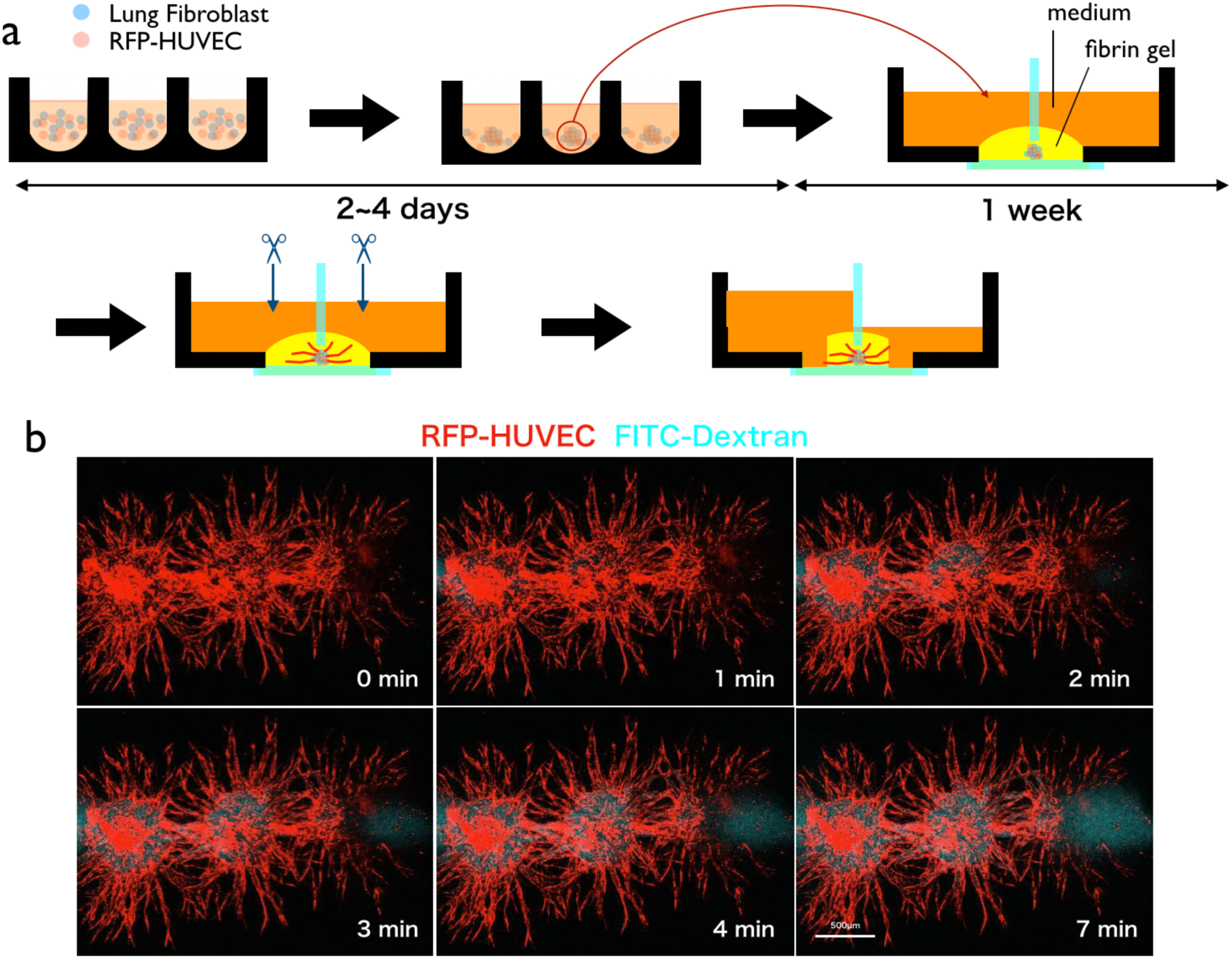
Perfusion of vascularized spheroids. (a) Experimental procedure. Spheroids containing RFP-HUVECs and lung fibroblast were generated and embedded in fibrin gel. After 1 week, sprouts from the spheroids became sufficiently long. Then, we cut the tip of the sprouts from both sides of the well and exerted static pressure to one side of the well. (b) Visualization of the perfusion inside the spheroid using FITC-dextran. Scale bar: 500 µm.

By exerting static pressure between two wells, we observed the medium flow in the connected spheroids (Fig. 8b). Interestingly, if we set two spheroids in close vicinity, the vascular sprouts from both spheroids fused and made a serial cluster of spheroids with perfusable vasculature.

## Discussion

### Reconstructive system of pattern formation

In the present study, we constructed a simple perfusion system to reproduce the flow in a capillary network. The seminal work on this type of culture system with flow was performed by Noo-Li Jeon’s group [14]. However, the system needs an elaborate microfluidic device that can only be fabricated by researchers with access to clean room facilities, and it is time consuming for biologists to fully use the system including the device, tubing, and syringe pump. In contrast, our system employs a standard glass-bottom dish without any additional pump, tubing, reservoir, or engineering. The simplicity of the system enabled us to combine it with time-lapse observation on a stage top incubator without any additional apparatus. However, there are several shortcomings because of its simplicity. For example, we could not regulate the flow rate in this system accurately because the driving force for perfusion is static pressure between two wells. For detailed control of flow, we need to use microfluidic devices.

### Lumen formation capability of endothelial cells

The mechanism of lumen formation by endothelial cells remains to be elucidated. ECM degradation and apicobasal polarity formation are thought to play a role [27]. In our culture system, one factor was the diffusible signaling molecules from LFs. HUVECs cultivated in fibrin gel formed a network even without LFs, but we could not induce a lumen without coculture with LFs. Two reports used a proteomics approach to identify the diffusible factor responsible for lumen induction by LFs. They identified several factors, but the effect of the combination of these factors was less effective than lung fibroblast coculture [28, 29]. We also found that lumens were preferentially formed at regions near a hard object (Fig. S2). One possible mechanism may be high cell density due to preferential migration of cells toward the hard substrate (durotaxis) [30], but increasing the cell density as a whole did not increase the lumen formation region around hard objects. In addition, the lumen formation capacity of HUVECs and LFs differed significantly depending on the production batch. We sometimes experienced HUVECs and LFs whose proliferation appeared to be normal, but lumen formation was poor. Increasing cell numbers sometimes compensated for the low lumen formation activity, but in many cases, using a different batch of cells improved lumen formation.

### Effect of the extracellular matrix on the pattern formation mechanism

We used fibrin gel to generate a perfusable vascular network based on a previous study [14]. Fibrin is the main component of a blood clot and not used in physiological tissues. We tried to substitute fibrin gel with other physiological extracellular matrixes such as Matrigel and type I collagen without success. This may be due to the physical viscosity difference. A fibrin gel solution is less viscous than a collagen gel solution. As a result, suspended HUVECs in fibrin gel gathered at the bottom of a dish because of gravity. We succeeded in generating a lumen structure in a type I collagen gel by embedding HUVEC spheroids, indicating that the local cell density may be critically important.

### Dynamics of pericytes in the capillary network with flow

For recruitment of pericytes to nascent blood vessels, platelet-derived growth factor B (PDGF-B) produced by tip endothelial cells plays pivotal roles [25, 31]. Notably, tip endothelial cells adjacent to hypoxic tissues are exposed to high concentrations of VEGF-A [31]. Moreover, because of the absence of lumens, tip endothelial cells have limited access to circulating blood [31]. Consistently, cultured HUVECs under high oxygen pressure exhibit reduced production of PDGF-B [32]. In our pericyte-HUVEC coculture with flow, pericytes unexpectedly disappeared in the flow region. A possible reason may be the lack of PDGF-B in the flow region. Endothelial cells in the flow region were under normoxic conditions. There are no flow regions in which endothelial cells should produce PDGF-B [32], and because of the PDGF-B gradient, pericytes might migrate away from the flow region or extensively proliferate at non-flow regions.

### Connection of two vasculature systems

The connection between two different capillary systems still remains a major challenge. We tried to establish connections between the self-organized HUVEC vascular network and the vascular network in embryonic tissue. The endothelial cells from the embryonic tissue appeared to adhere to the HUVEC structure. However, the lumens of these vasculatures did not form a connection. Therefore, we could not perfuse the vasculature in embryonic tissue (Fig. S6). Additional factors may be necessary to use this culture system as an alternative to organ culture. Because direct connection of two vascular systems is still a major challenge, currently, a technically simpler experimental model of vasculature-blood cell interactions may be a major field of application. In the present study, we demonstrated two applications, dynamics of inflammatory cells and hematogenic metastasis. Various other system can be implemented with our current model without additional costs.

### Relationship between microfluidic device methods

This method may be a very good prescreening method for microfluidic device experiments. The major merit of this method is the ease and low cost. We have experienced difficulty in introducing blood vessel into a microfluidic device [15] because it requires specialized skills to connect tubes without damaging gels or endothelial cell networks. As a result, it is difficult to perform a large number of microfluidic device experiments because of its technical difficulty. Although we only used a constant pressure condition for medium perfusion and could not strictly control the flow, this method may be a good starting point to roughly estimate the effect of flow in biological laboratories.

## Acknowledgments

The authors thank Professor Fumio Arai for helpful suggestions. We also thank Mitchell Arico from Edanz Group (https://en-author-services.edanzgroup.com/) for editing a draft of this manuscript. This work was financially supported by JST CREST (Grant Number JPMJCR14W4).

## Author contributions

Kei Sugihara, Yoshimi Yamaguchi, and Shiori Usui undertook experiments. Yuji Nashimoto and Sanshiro Hanada provided materials and designed experiments. Etsuko Kiyokawa provided tumor cell lines. Akiyoshi Uemura designed experiments, provided materials, and wrote the pericyte-related section in the manuscript. Ryuji Yokokawa, Koichi Nishiyama, and Takashi Miura designed experiments and wrote the manuscript.

## Supporting information

Supporting movie S1: Flow of fluorescent particles in the self-organized capillary network. Real-time movie of fluorescent particles flowing in a synthetic vascular network in fibrin gel.

Supporting movie S2: Flow of red blood cells in the self-organized capillary network. Real-time movie of red blood cell flowing in a synthetic vascular network in fibrin gel. Blood was diluted 20× using EGM-2 and loaded on one side of the well (Fig. 1k,l).

Supporting movie S3: Cell debris flowing during the medium change (long-term perfusion experiment). Real-time movie of cell debris flowing in a synthetic vascular network in fibrin gel during the long-term perfusion experiment (Fig. 2). We confirmed perfusion during the medium change process.

Supporting movie S4: dilation of the vessels in the flow region Time-lapse movie of the flow region in the long-term flow experiment (Fig. 3d). The vessel with flow was maintained alive, and some vessels became dilated. Frame rate: 1 day/frame, 30 days.

Supporting movie S5: Disappearance of unused vessels in the non-flow region. Time-lapse movie of the non-flow region in the long-term flow experiment (Fig. 3e). The vessel became disconnected and degraded gradually. Frame rate: 1 day/frame, 30 days.

Supporting movie S6: Movement of pericytes in the flow region Time-lapse movie of pericytes (green) in the flow region (upper half) and non-flow region (lower half). In the flow region, the pericytes simply disappeared, whereas in the non-flow region, pericytes proliferated extensively.

Supporting movie S7: Flow of HL60 cells in the self-organized vascular network. Time-lapse movie of HL60 (green) in HUVEC-RFP vascular network.

---

**S1 Fig:**
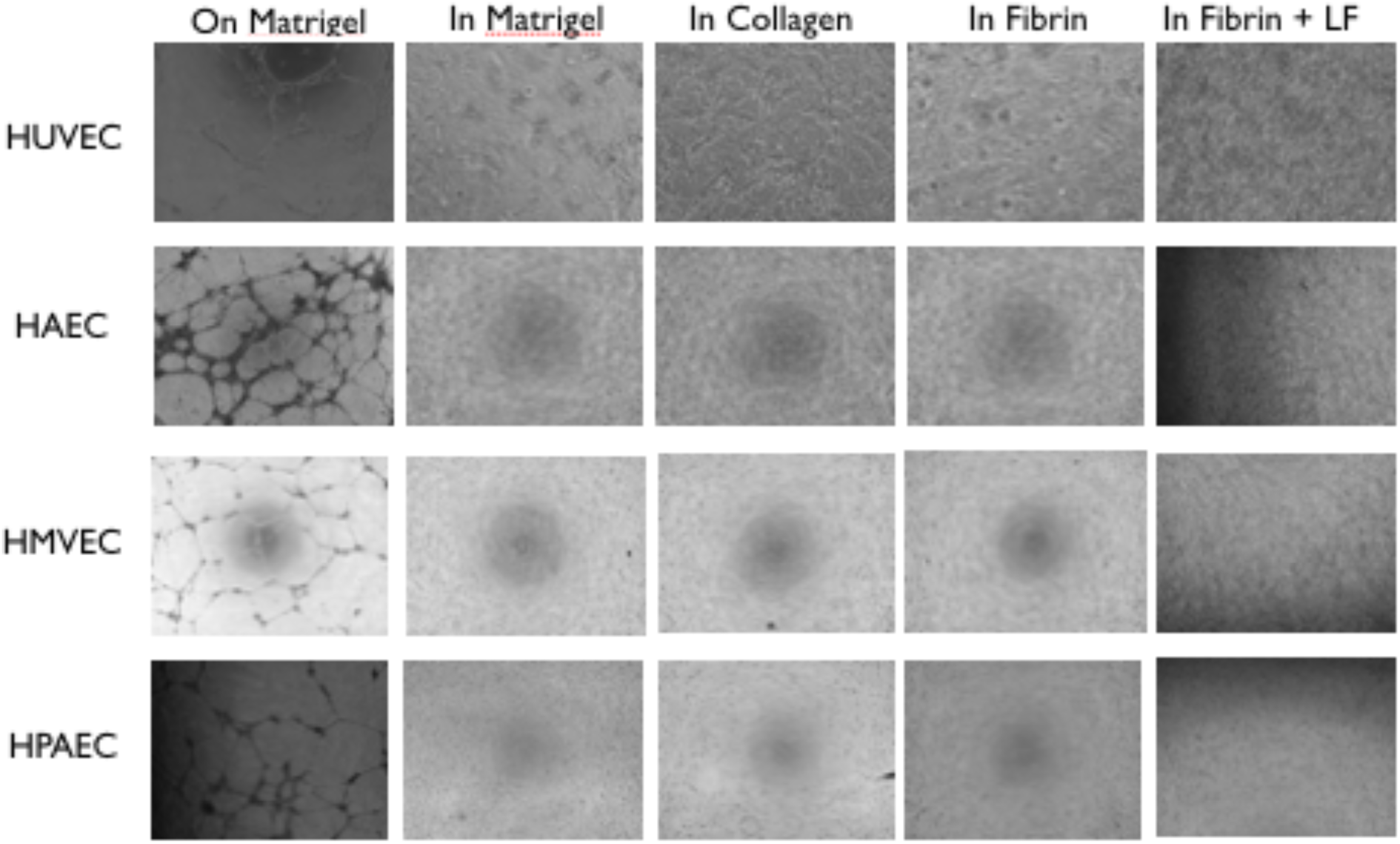
Screening for the pattern formation capability of various commercially available primary endothelial cells. All cells generated a meshwork structure on Matrigel (tube formation assay). In Matrigel or collagen gels, the cells simply became static and no pattern formation phenomena was observed. In fibrin gel, the cells tended to connect to each other, but a lumen was not formed, except for HPAECs. If we cocultured these cells with human lung fibroblasts (LFs), the cells formed a network with lumens.

**S2 Fig:**
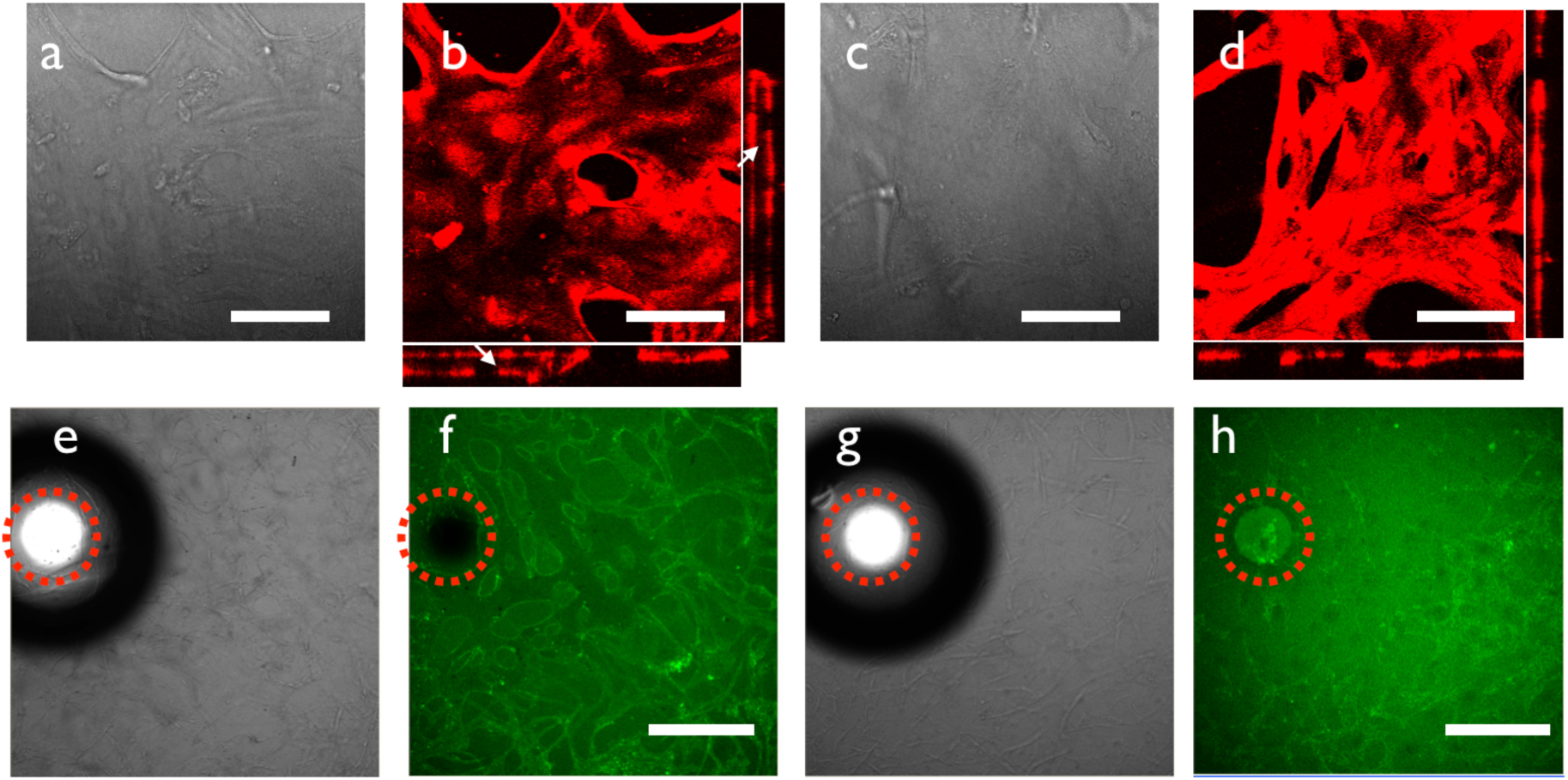
Lumen formation was observed only around solid objects. (a) Brightfield image of HUVECs cocultured with LFs in fibrin gel in the *periphery* of the dish. A lumen structure was observed. (b) Fluorescence image of HUVECs cocultured with LFs in fibrin gel in the *periphery* of the dish. HUVECs were stained with UEA1-FITC. A lumen structure was observed (white arrows). (c) Brightfield image of HUVECs cocultured with LFs in fibrin gel in the *center* of the dish. A lumen structure was not clear. (d) Fluorescence image of HUVECs cocultured with LFs in fibrin gel in the *center* of the dish. HUVECs were stained with UEA1-FITC. Lumen formation was not clear. (e) Brightfield image of HUVECs cocultured with LFs in fibrin gel near 1 mm glass beads *embedded in* the fibrin gel. We observed lumen formation around beads. (f) Confocal image of HUVECs cocultured with LFs in fibrin gel near 1 mm glass beads (dashed red circle) *embedded in* the fibrin gel. HUVECs were stained with UEA1-FITC. We observed lumen formation around beads. (g) Brightfield image of HUVECs cocultured with LFs in fibrin gel near 1 mm glass beads *placed on* the fibrin gel. We observed lumen formation around beads. (h) Confocal image of HUVECs cocultured with LFs in fibrin gel near 1 mm glass beads *placed on* the fibrin gel. HUVECs were stained with UEA1-FITC. We observed lumen formation around the beads. Scale bars: 100 µm (a‒d); 1 mm (f, h).

**S3 Fig:**
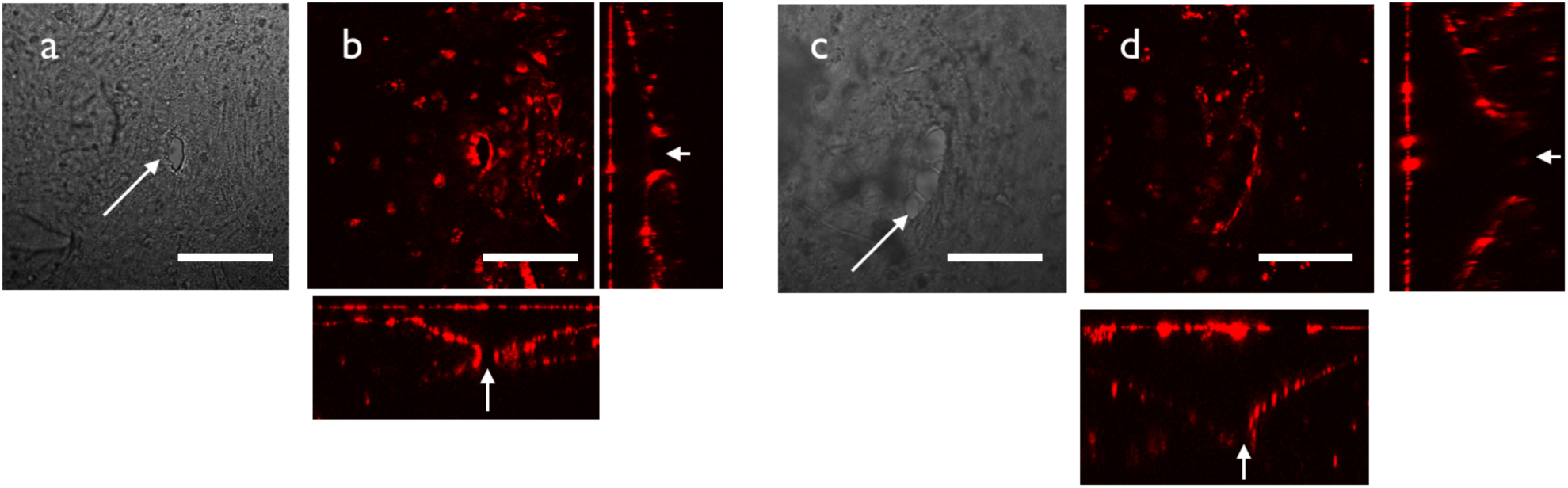
Three-dimensional structure of inlet and outlet regions. (a) Brightfield image of the inlet hole region. Arrow: small hole made by fine forceps. (b) Three-dimensional structure of the inlet hole observed by confocal microscopy. We observe RFP-HUVECs covering the hole to make openings to the upper medium reservoir. (c) Brightfield image of the outlet hole region. Arrows indicate a small hole made by fine forceps. Cell debris from the vascular network accumulated near the outlet region (arrowhead). Scale bars: 200 µm.

**S4 Fig:**
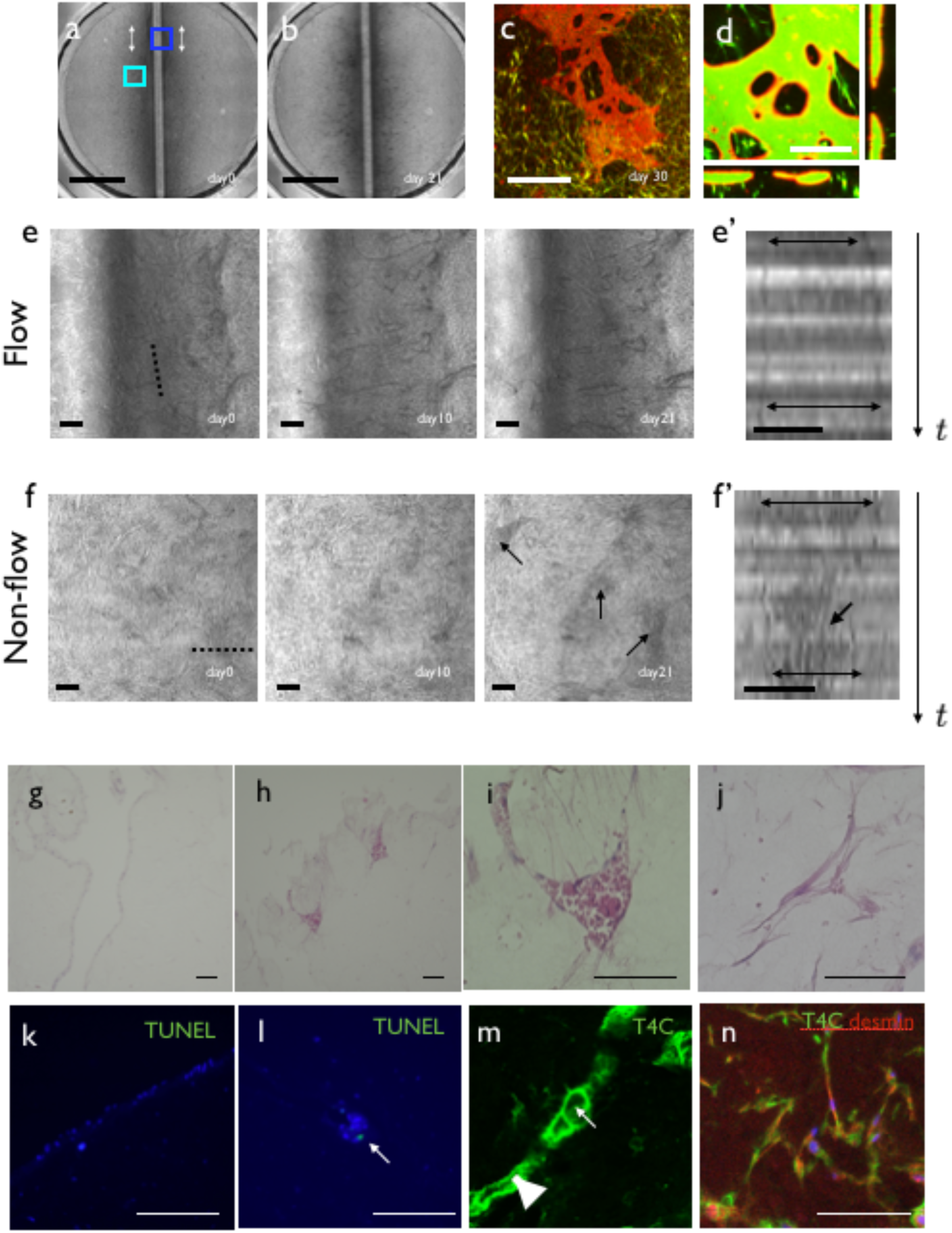
Effect of flow on the self-organized endothelial cell network with pericytes. (a) Low magnification view at day 0. Locations of the inlet and outlet are shown by white arrows. (b) Low magnification view at day 21. (c) Confirmation of perfusion at day 30. Endothelial cells were stained with UEA-1 lectin (red) and culture medium with FITC-dextran (green) was perfused. Characteristics of the vessel shape were similar to that without pericytes. (d) Three-dimensional view of the vasculature. We did not observe leakage of FITC-dextran. (e) High magnification view of the flow region [blue box in (a)]. Note that vascular regions increased gradually. (e’) Kymograph of the vascular region [dotted line in (e)]. Note that the vascular diameter increased gradually (double-headed arrows). (f). High magnification view of the non-flow region [cyan box in (a)]. The vascular region decreased gradually with cell debris inside the vascular lumen (double-headed arrows). (f’) Kymograph of vascular region [dotted line in (f)]. Vascular diameter decreased gradually and debris accumulated inside the endothelial cyst (double-headed arrows). (g) Histological observation of the flow region. (h) Histological observation of the low flow region. (i) High magnification view of the cyst structure in the non-flow region. (j) Histological structure of pericytes within a gel in the non-flow region. (k) TUNEL staining of the flow region. No dead cells were observed. (l) TUNEL staining of the non-flow region. A positive signal was observed within the cyst. (m) Type IV collagen staining near the cyst. A large ECM sheath structure was observed (arrowhead) near the cyst (arrow). (n) Type IV collagen and desmin staining of the non-flow region. Desmin-positive pericytes were observed within the gel, which colocalized with type IV collagen. Scale bars: 3 mm (a, b); 1 mm (c); 250 µm (d);100 µm (e, f).

**S5 Fig:**
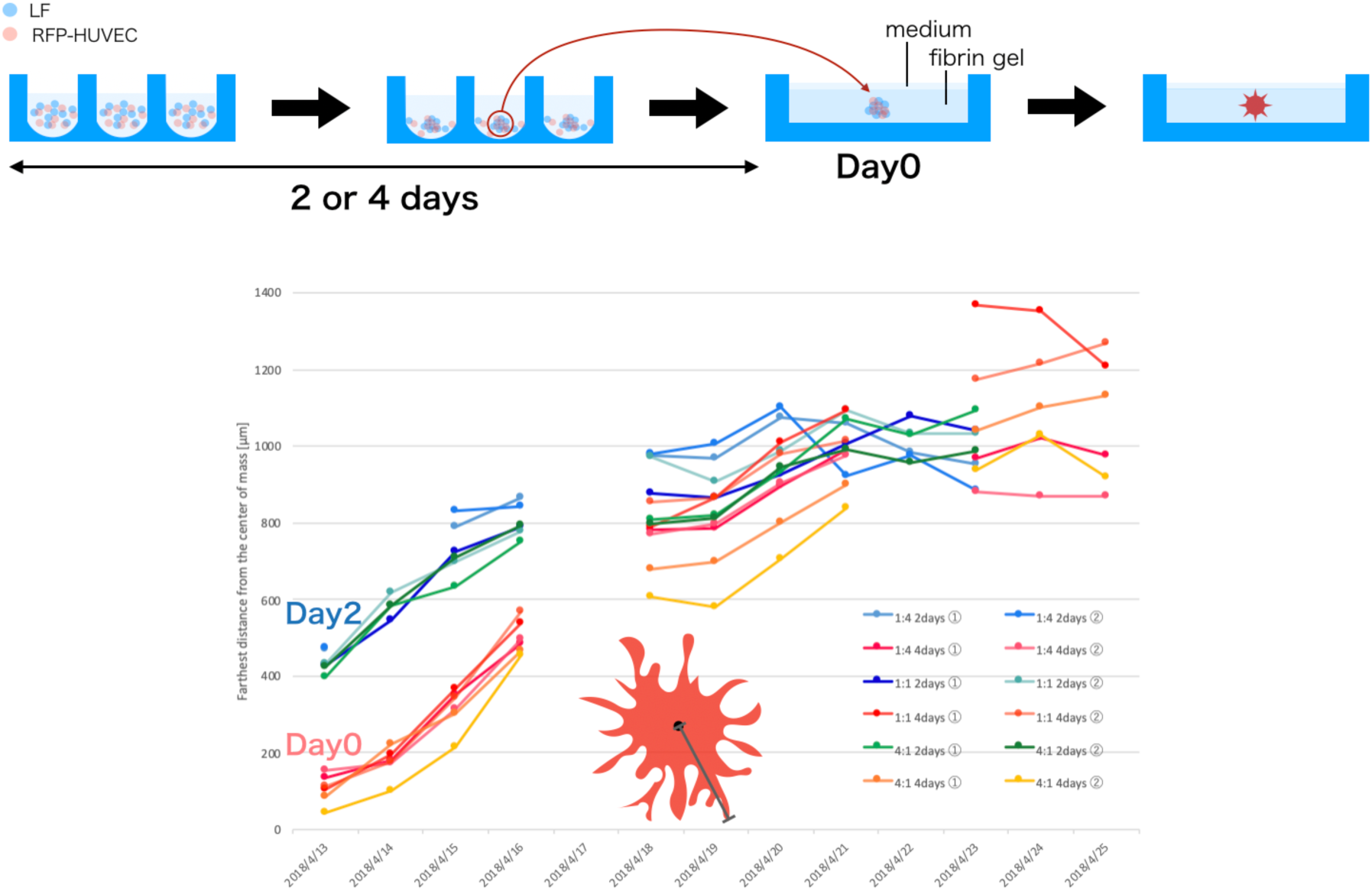
Quantification of the sprout length from endothelial cell-containing spheroids. Spheroids containing RFP-HUVECs were cultivated for 2 or 4 days in a culture plate and then transferred to fibrin gel. The length of angiogenic sprouts from the centroid of the fluorescent signal was measured every day. The length of the sprout became saturated after 7 days, and the radius was around 1 mm, which enabled us to directly cut the tip of the sprout to generate an open end.

**S6 Fig:**
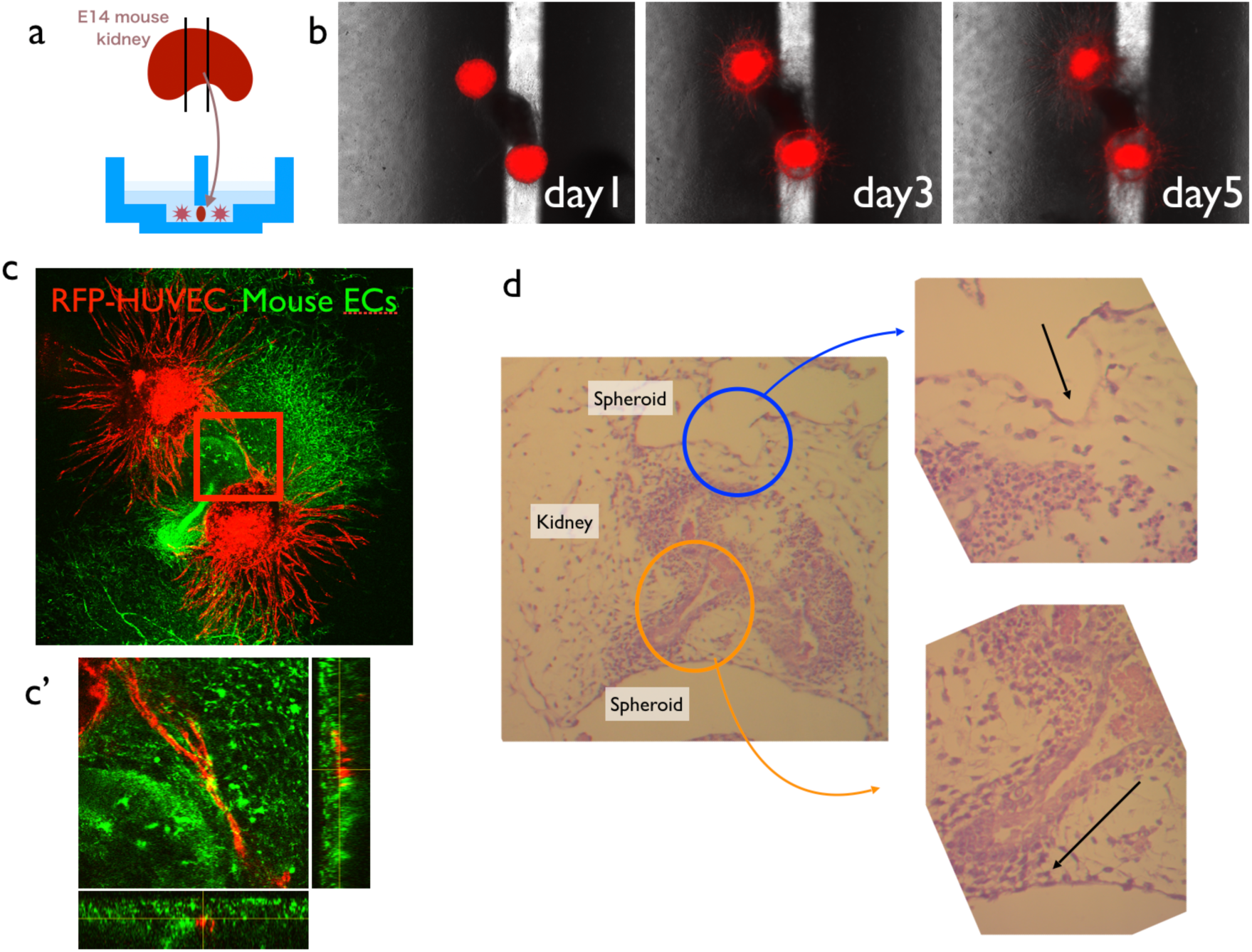
Attempt to connect vasculature to the embryonic tissue. (a) E12 mouse embryonic kidney tissue was dissected, and the central one-third of the kidney was embedded in fibrin gel. The tissue was sandwiched by two RFP-HUVEC:hLF spheroids. (b) After 1 week, HUVEC-LF spheroids generated sprouts towards the embryonic kidney tissue. (c) Sprouts from RFP-HUVECs appeared to avoid the embryonic tissue. (c’) 3D observation of HUVECs and mouse endothelial cells revealed that, although mouse endothelial cells (green) tended to attach to RFP-HUVECs, they did not connect vasculature with lumens together. (d) Histological observation of the cultured tissue. Kidney explants formed collecting tubule-like structures, but the endothelial sprouts from the HUVEC spheroids did not form a connection with the kidney structure.

